# CD161 mediates prenatal immune suppression of IFNγ-producing PLZF^+^ T cells

**DOI:** 10.1101/305128

**Authors:** Joanna Halkias, Elze Rackaityte, Dvir Aran, Ventura F. Mendoza, Walter L. Eckalbar, Trevor Burt

## Abstract

While the fetal immune system defaults to a program of tolerance, there is concurrent need for protective immunity to meet the antigenic challenges after birth. Activation of fetal T cells is associated with fetal inflammation and the termination of pregnancy, yet which fetal T cells contribute to this process is poorly understood. Here we show a transcriptionally distinct population of pro-inflammatory T cells that predominates in the human fetal intestine. Activation of PLZF^+^ T cells results in rapid production of Th1 cytokines and is inhibited upon ligation of surface CD161. This mechanism of fetal immune suppression may inform how immune dysregulation could result in fetal and neonatal inflammatory pathologies such as preterm birth. Our data support that human development of protective adaptive immunity originates *in utero* within the specialized microenvironment of the fetal intestine.

## INTRODUCTION

The developing human immune system is uniquely adapted for fetal survival within the semi-allogeneic environment *in utero* as well as for neonatal protection after contact with the many antigenic challenges encountered after birth. Unlike mice, which lack peripheral T cells *in utero*, human T cells begin to populate peripheral organs (including mucosal sites) by 11-14 weeks of gestation ^1, 2^. While naïve fetal T cells preferentially generate induced regulatory T cells upon antigen encounter in the periphery ^3^, there is evidence of protective fetal adaptive immunity in response to pathogens and vaccines ^4–6^. However, activated T cells are associated with the fetal inflammatory response and contribute to the comorbidities associated with preterm birth ^7–11^. The concurrent development of fetal tolerogenic and protective T cell programs indicate that human immune development may be more nuanced than previously appreciated.

The presence of functional memory T cells in the fetus and infant ^12–15^ indicate that adaptive immune memory originates *in utero*. However, the spatial compartmentalization of human T cell differentiation and function suggests that the effector capacity of the fetal immune response cannot be inferred from blood ^13,16,17^. The existence of organized intestinal lymphoid structures as early as the second trimester of gestation ^18^, along with the environmental, maternal, and selfantigens within swallowed amniotic fluid, points to an instructive role for the intestinal mucosa in the development of fetal adaptive immunity.

In this study, we performed a detailed examination of fetal lymphoid and mucosal TCRαβ CD4^+^ T cells. We identified a polyclonal population of IFNγ-producing T cells characterized by expression of the transcription factor Promyelocytic Leukemia Zinc Finger (PLZF) and the C-type lectin CD161. These Va7.2^-^ PLZF^+^ CD161^+^ TCRαβ^+^ CD4^+^ T cells preferentially accumulate in the lamina propria (LP) of the small intestine and possess a resident memory phenotype inferred from expression of CD69 and CD103 ^19^. We demonstrate that Va7.2^-^ PLZF^+^ CD161^+^ TCRαβ+ CD4^+^ T cells, herein referred to as polyclonal intestinal (pi) PLZF^+^ CD161^+^ T cells, are transcriptionally distinct and possess a type 1 T helper (Th1) functional signature, which identifies them as a major effector population in the fetal intestine. We also show that the ability of these fetal T cells to produce IFNγ is positively associated with advancing gestation. We found that Lectin-like transcript 1 (LLT1), the ligand for CD161, was uniquely expressed by resident macrophages of the intestine, and that ligation of CD161 inhibited the TCR-mediated activation of pi-PLZF^+^ CD161^+^ T cells. We propose a model in which functional maturation of pi-PLZF^+^ CD161^+^ T cells occurs under steady state in a spatially segregated manner in the lamina propria of the intestine, where CD161 mediates immune suppression and allows for induction of immunity uncoupled from inflammation. The ability of pi-PLZF^+^ CD161^+^ T cells to respond to both TCR- and cytokine-mediated signaling suggests a potential for these cells to contribute to the multitude of inflammatory pathologies associated with premature birth. Our identification of the suppressive role of CD161 points to a putative therapeutic target to control the fetal inflammatory response for which there is currently no effective therapy.

## RESULTS

### Memory PLZF^+^ CD4^+^ T cells are highly enriched in the fetal small intestine

The human fetal thymus produces a subset of CD4^+^ T cells which express the transcription factor PLZF and are distinct from iNKT cells and MAIT cells1 ^20^. To explore whether PLZF^+^ T cells contribute to the development of protective immunity *in utero*, we examined their distribution in mucosal and lymphoid tissue. PLZF expression was abundant among lamina propria (LP) CD4^+^ T cells of the fetal small intestine (Fig. 1a). As many human innate cells express PLZF (cite), we restricted our analysis to TCRβ^+^ Va7.2^-^ CD4^+^ T cells, which excluded innate lymphoid cells, MAIT cells, and γδ T cells (Supplementary Fig. 1). PLZF^+^ T cells accounted for ~30% of all intestinal Va7.2^-^ TCRαβ+ CD4^+^ T cells, whereas they averaged <10% of CD4^+^ T cells across all other lymphoid and non-lymphoid sites with the exception of the appendix (Fig. 1b, c). Adult human PLZF^+^ T cells often co-express the C-type lectin CD161, which identifies T cells with innate characteristics ^21^. The majority of intestinal PLZF^+^ T cells and ~ half of those in the appendix were CD161^+^, whereas less than half of PLZF^+^ T cells across other sites expressed CD161 (Fig. 1b, d). Consistent with the reported decline in thymic production of PLZF^+^ CD4^+^ T cells with advancing gestation ^20,22^, PLZF^+^ CD4^+^ T cells are essentially undetectable in term cord blood (CB) and adult peripheral blood (aPB), and we used aPB samples as internal controls between experiments (Extended Fig. 1a-b).

**Figure 1:**
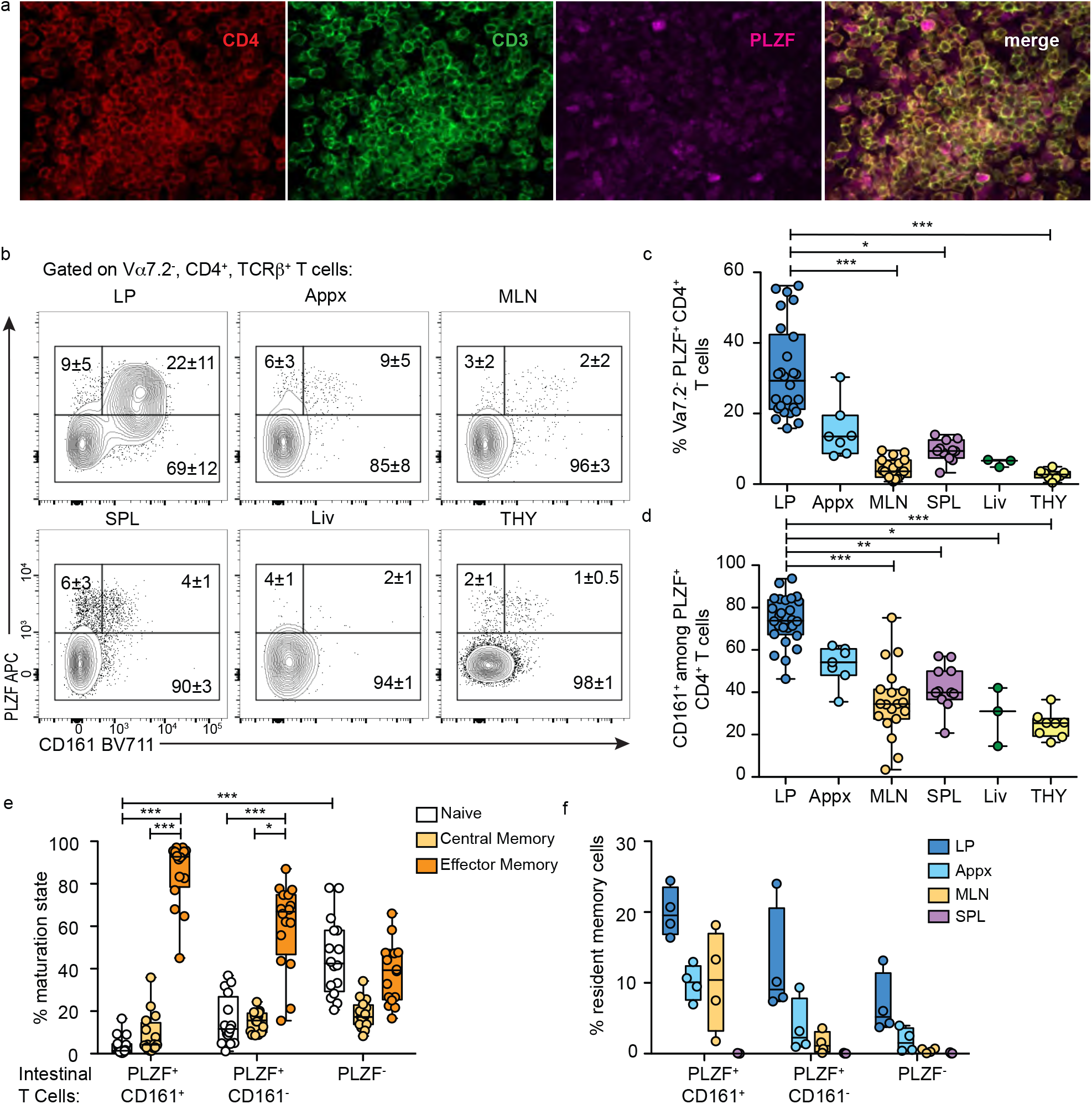
The fetal small intestine is enriched for PLZF+ CD4+ T cells. a, Representative imaging (40×) of PLZF expression in the lamina propria of the small intestine (n=3). b, Representative flow plots of PLZF and CD161 expression among CD4+ T cells in fetal tissues. c, Frequencies of PLZF+ CD4+ T cells and d, Proportion of CD161+ cells among PLZF+ CD4+ T cells in fetal tissues. e, Frequencies of Naïve (CD45RA+, CCR7+), Central Memory (CD45RA-, CCR7+), and Effector Memory (CD45RA-, CCR7-) phenotypes within indicated populations of intestinal CD4+ T cells. f, Frequencies of resident memory populations within indicated subsets of CD4+ T cells. Circles represent individual donors (c-d, e-f). Significance was measured by Kruskal Wallis paired ANOVA with Dunn’s multiple comparison test (c-d, e-f). *p < 0.05, **p < 0.005, ***p < 0.0005. Numbers in the flow cytometry plots correspond to the frequencies of gated populations ± SD.

Previous studies have identified memory T cells in fetal and neonatal small intestine ^12,13,23,24^, which led us to examine the maturation state of intestinal T cells on the basis of CD45RA and CCR7 expression. The majority of intestinal PLZF^+^ CD4^+^ T cells had a CD45RA^-^ CCR7^-^ effector memory (T_EM_) phenotype (85% of PLZF^+^CD161^+^ and 60% of PLZF^+^ CD161^-^) and contained significantly less naïve cells than PLZF^-^ T cells (Extended Fig. 1c & Fig. 1e). This held true across multiple tissues, where PLZF^+^ CD161^+^ T cells were significantly more likely than PLZF^-^ T cells to possess a T_EM_ phenotype in the MLN, and trended towards more T_EM_ in the appendix and spleen (Extended Fig. 1d). In contrast, PLZF^-^ CD4^+^ T cells had a more even distribution of naïve, central memory (T_CM_) and T_EM_ cells, similar to that reported for infant and pediatric intestine ^12,13^. Most intestinal PLZF^+^ T cells were CD69^+^, and ~20% also expressed CD103 consistent with a resident memory (T_RM_) phenotype (Extended Figure 2e) ^13,16,25^. Overall, a T_RM_ phenotype was most frequent among intestinal PLZF^+^CD161^+^ T cells (Fig. 1f).

**Figure 2:**
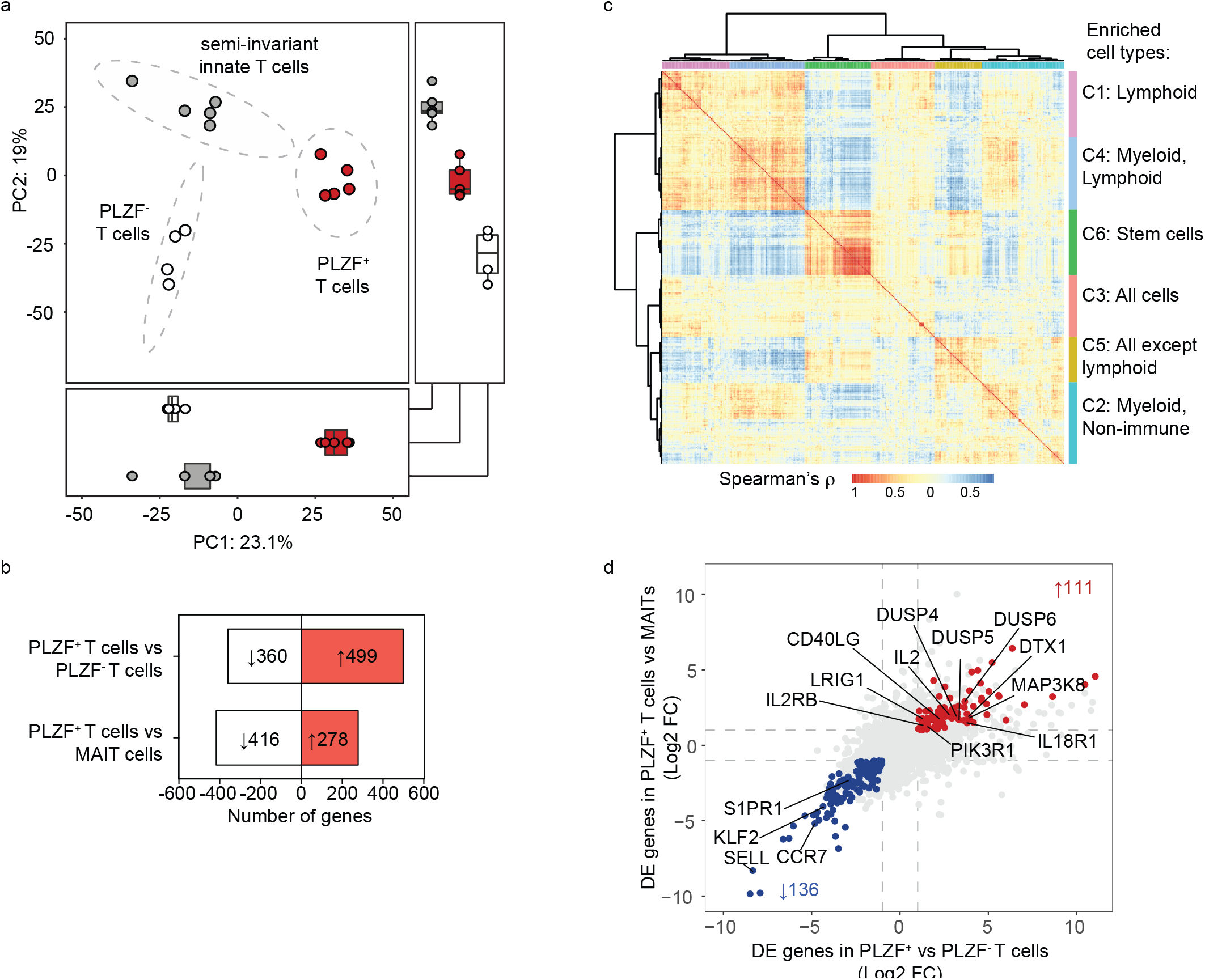
PLZF+ CD161+ CD4+ T cells are a distinct population of fetal T cells. a, Principal component analysis (PCA) of the top 1000 variable genes identified by RNAseq of fetal intestinal PLZF^+^ T cells (n=5), PLZF^-^ T cells (n=4), and semi-invariant innate T cells cells n=5). Ellipses denote the 95% confidence intervals of the population means (PERMANOVA R2 0.92, p<0.001). Boxplots indicate scores for PC1 (bottom), and PC2 (right). b, Numbers of genes with >2 fold (FDR<0.05) increase (red) or decrease (white) in expression levels between the indicated populations. c, Correlation analysis using the Human Primary Cell Atlas as a reference of the differentially expressed genes (>2 fold, FDR<0.05) in PLZF+ T cells as compared to PLZF-T cells revealing clusters of genes (C1-C6) that are co-expressed in the same cell types. d, Multi-way comparison of differentially expressed genes (>2 fold, FDR<0.05) in PLZF+ T cells compared to MAIT cells (y-axis) and compared to PLZF- T cells (x-axis) identifies a core signature of genes uniquely enriched (red) and uniquely depleted (blue) within PLZF+ T cells. Selected genes involved in immune response, immune regulation, and leukocyte migration are labeled.

### PLZF^+^ CD161^+^ CD4^+^ T cells are a transcriptionally distinct population of mucosal T cells

To investigate whether intestinal PLZF^+^ T cells represent a distinct population of fetal T cells, we determined their global gene expression profile and examined their relationship to innate, semiinvariant Va7.2^+^ T cells and to conventional PLZF^-^ CD4^+^ memory T cells (Extended Fig. 2a). We sorted semi-invariant innate T cells on the basis of TCRβ^+^ Vα7.2^+^ CD161^+^ cells ^26,27^, of which the majority (>75%) were also IL18R+ (Extended Fig. 2b). Although not exclusive for mucosal-associated invariant T (MAIT) cells, this subset is enriched for MAIT cells ^28^. We identified additional surface markers that allowed us to separate PLZF^+^ from PLZF^-^ memory CD4^+^ T cells (Extended Fig. 2b,c, Supplementary Table 1). The expression of *ZBTB16, IL18R, PDCD1*, and *KLRB1* (the respective genes for PLZF, IL18R, PD1 and CD161) mirrored their protein expression and validated our sorted T cell populations and our gene expression analysis (Extended Fig. 2d). Intestinal PLZF^+^ T cells formed a spatially separate group from either semiinvariant innate T cells or PLZF^-^ memory T cells, reflecting a divergent transcriptional state by principal component analysis (PCA) (Fig. 2a). PLZF^+^ T cells were highly distinct by principal component (PC) 1 (23% of variance) and segregated between PLZF^-^ T cells and semi-invariant innate T cells along PC2 (19.9% of variance), suggesting shared attributes with these populations. Twice as many genes were enriched in PLZF^+^ relative to PLZF^-^ memory T cells (499) as compared to semi-invariant innate T cells (278) (>2-fold, FDR<0.05)(Fig. 2b).

We first focused on the genes that differentiated PLZF^+^ T cells from conventional PLZF^-^ T cells and utilized the Human Protein Cell Atlas as a reference to identify genes that cluster together by cell type using correlation analysis ^29^. This identified 6 clusters, of which clusters 2 and 4 were comprised predominantly of myeloid-associated genes (Fig. 2c, Extended Fig. 3a). Among the myeloid genes in cluster 4 were those associated with immune regulation (GAB2, NLRP3, TGFB1, and C3AR1), while myeloid-enriched genes in cluster 2 were involved in lipid metabolism (PPARG, PLIN2, and NFIL3) (Extended Fig 3b). Cluster 6 was enriched for stem cells and contained genes involved in the cell cycle from which we inferred that PLZF^+^ T cells were likely proliferating *in vivo* (Extended Fig. 3b, Supplementary Figure 2). Cluster 1 was enriched for genes associated with immune activation and overlapped with effector memory T cells, NK cells, and γδ T cells suggestive of effector properties including Th1-like and cytotoxic functions (Extended Fig. 3b, Supplementary Figure 2). This correlation analysis allowed us to extrapolate function based on expression overlap with other cell types, and revealed an atypical composition of myeloid-, lymphoid-, and stem cell-associated genes in PLZF^+^ T cells. A striking difference in the genes differentially expressed between PLZF^+^ T cells and semiinvariant innate T cells was the diversity of TCR usage. A number of TRAV regions were enriched in PLZF^+^ T cells, while semi-invariant innate T cells displayed preferential expression of TRBV9, TRBV7-9, and TRBV6-4 which have been associated with MAIT cells ^30,31^ (Extended Fig. 3d). The polyclonal nature of PLZF^+^ T cells was supported by variable Vβ use among both PLZF^+^ and PLZF^-^ T cells and was comparable to that reported in fetal blood ^32^ with no single chain accounting for >10% of TCR Vβ families (Extended Fig. 3e).

**Figure 3:**
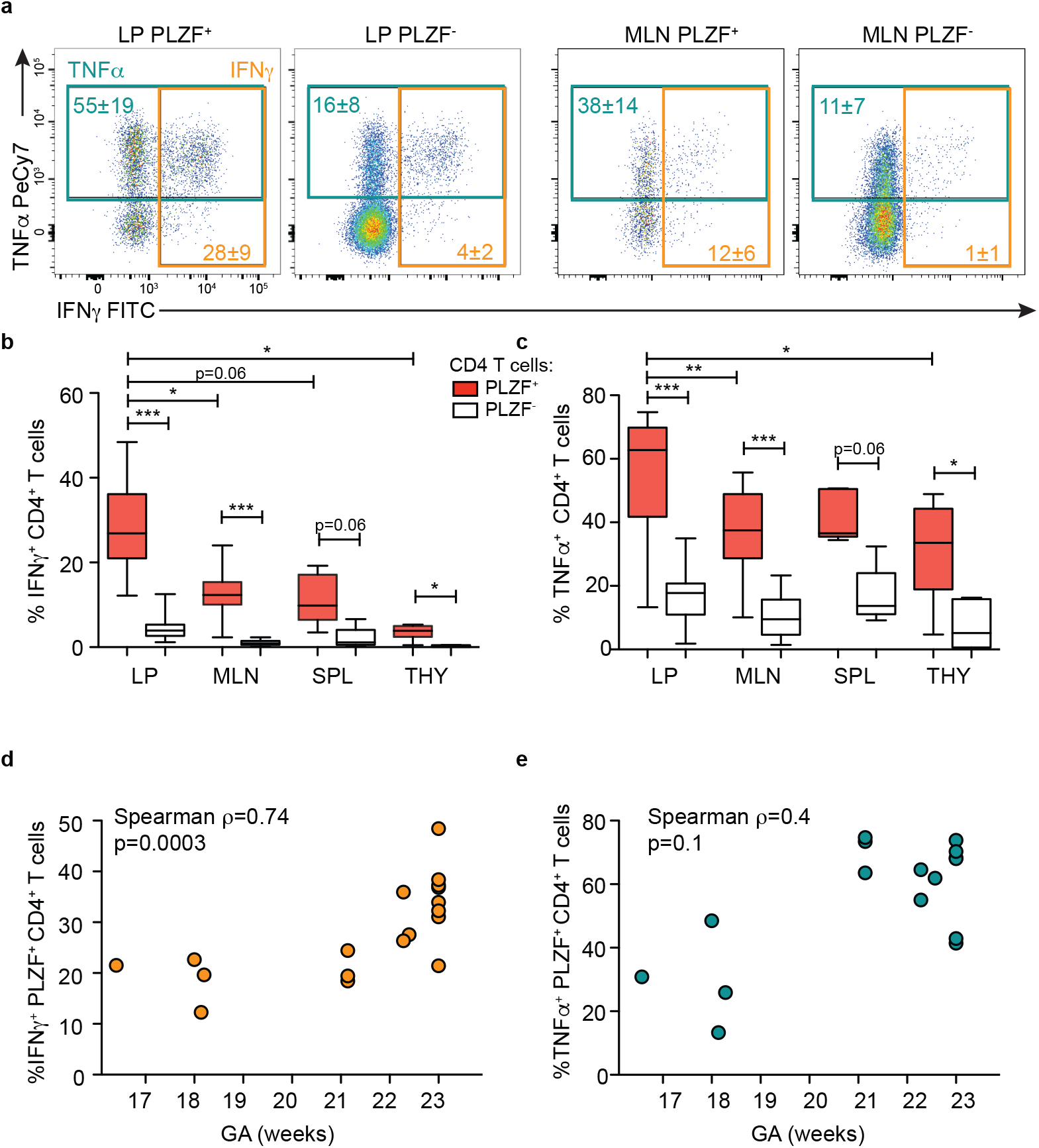
PLZF+ CD4 T cells possess a distinct functional signature. a, Representative flow plots of intracellular TNFα and IFNγ staining of indicated populations of intestinal, MLN, and adult PB live Vα7.2-TCRβ+CD4+T cells after PMA/Ionomycin stimulation. Numbers represent mean ± SD of gated frequencies. (n=18). Frequencies of b, IFNγ + and c, TNFα+ cells within indicated populations of Vα7.2-TCRβ+CD4+T cells after PMA/Ionomycin stimulation within fetal LP (n=18), MLN (n=11), SPL (n=5), and THY (n=6). d, e, Spearman’s rank correlation was used to calculate the association between d, IFNγ and e, TNFα production by intestinal PLZF+ CD4 T cells in response to PMA/Ionomycin and weeks of gestational age (GA). Circles represent individual donors. Wilcoxon matched-pairs signed rank test (b, c). *p < 0.05, ***p < 0.001.

We identified a core signature of 111 genes that were enriched, and 137 genes that were depleted in PLZF^+^ T cells compared to both PLZF^-^ T cells and semi-invariant innate T cells (>2FC; FDR<0.05) (Fig. 2d, Extended Fig. 3f). PLZF^+^ T cells demonstrated distinct gene expression profiles enriched in immune activation pathways, as well as genes involved in the regulation of the immune response (Fig. 2d). Specifically, the core signature of PLZF^+^ T cells contained transcripts involved in T cell activation (IL2, CD40LG, IL18R1, IL2RB, MAP3K8, PIK3R1), as well as T cell regulation (DUSP4, DUSP5, DUSP6, LRIG1, and DTX1). Among the genes specifically depleted in PLZF^+^ T cells were those required for lymph node homing and tissue egress (*SELL, CCR7, S1PR1, KLF2*)^19^, supporting the characterization of these cells as human T_RM_ cells. We therefore define a transcriptionally unique subset of polyclonal intestinal PLZF^+^ CD4 T cells, which share gene expression profiles with innate immune cells suggestive of a rapid effector phenotype. However, the presence of numerous negative T cell regulators among the gene signature of PLZF^+^ T cell points to cell-intrinsic mechanisms of regulation to promote immune homeostasis *in utero*.

### PLZF^+^ CD4 T cells possess Th-1 effector function

Lamina propria PLZF^+^ T cells produced large amounts of the Th1 cytokines TNFα and IFNγ upon short-term polyclonal activation by Phorbol 12-myristate 13-acetate (PMA) and Ionomycin, and expression of CD161 did not predict cytokine production among PLZF^+^ T cells (Fig. 3a, Extended Fig. 4a, b). Among CD4 T cells, IFNγ was predominantly produced by PLZF^+^ cells, and intestinal PLZF^+^ T cells produced more IFNγ than those of the mesenteric lymph node (MLN), spleen, or thymus (Fig. 3b). While PLZF^+^ T cells also made more TNFα than their PLZF^-^ counterparts, production of TNFα was less anatomically restricted and was already evident in >20% of thymic PLZF^+^ T cells (Fig. 3c). Further, the capacity of PLZF^+^ T cells to produce IFNγ was directly correlated with advancing gestation, but this association was not significant for TNFα, nor between either cytokine and PLZF^-^ T cells (Fig. 3d, e & Extended Fig. 4c, d). Consistent with the presence of *IL-2* in their core signature, intestinal PLZF^+^ T cells produced more IL-2 than PLZF^-^ T cells, however production of IL-2 was equally abundant among MLN T cells (Extended Fig. 4e,g). In contrast to Th1-asociated cytokines, IL-8 production was equivalent between PLZF^+^ and PLZF^-^ T cells and was lower in the intestine compared to the MLN (Extended Fig. 4f, h). Thus, in addition to a core transcriptional signature, PLZF^+^ T cells possess a Th1-type functional profile and are an abundant source of IFNγ in the fetal immune system.

**Figure 4:**
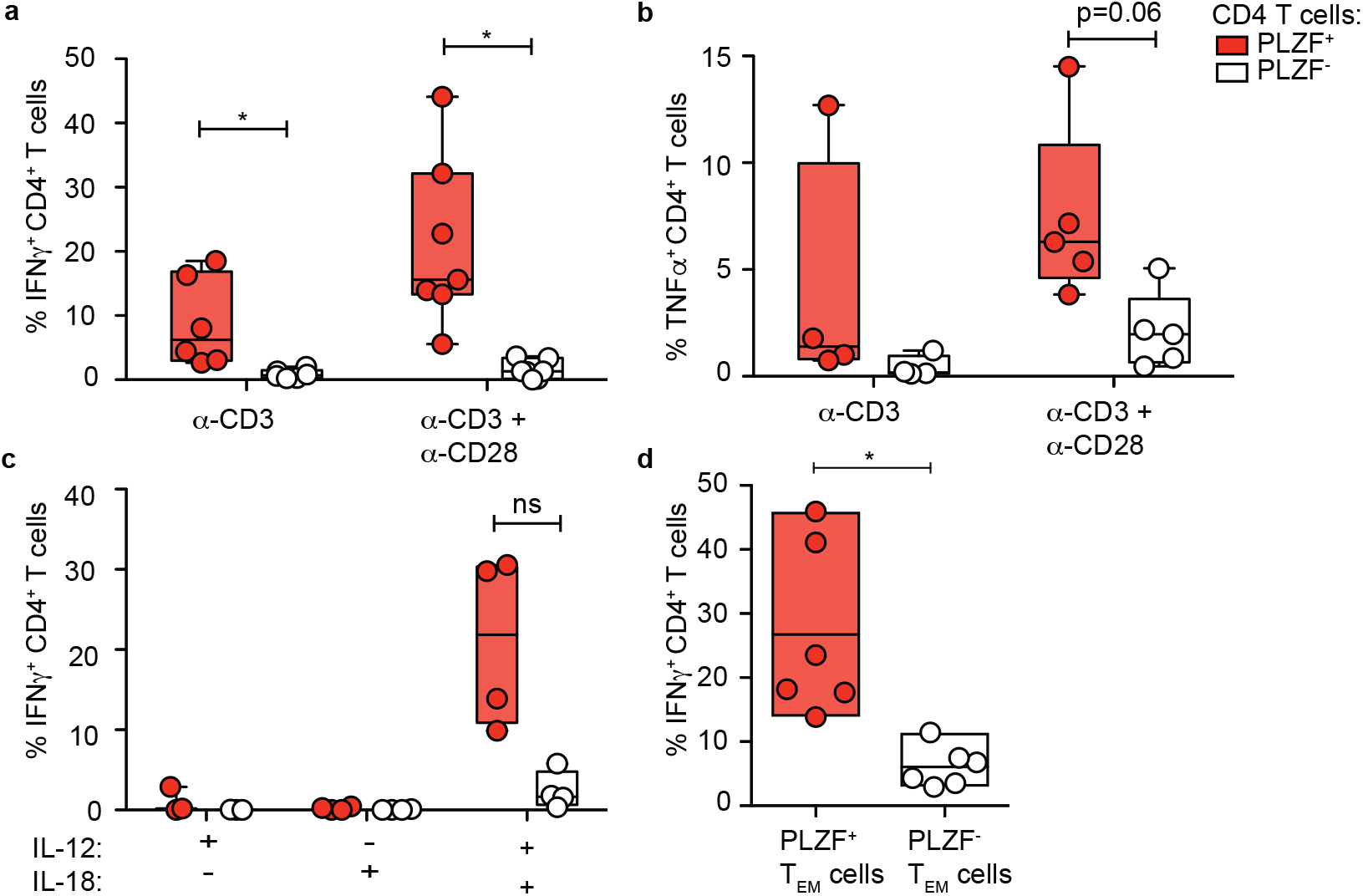
Dual activation of PLZF+ T cells. a, b Frequencies of a, IFNγ + and b, TNFα+ cells within indicated populations of intestinal live Vα7.2-TCRβ+CD4+T cells after 12-14 hrs of TCR stimulation with anti-CD3 +/- anti-CD28. c, Frequencies of IFNγ + cells within indicated populations of intestinal Vα7.2-TCRβ+CD4+T cells after 12-14 hrs of cytokine stimulation with IL-12 and/or IL-18. d, Frequencies of IFNγ + cells among intestinal PLZF+ and PLZF- TEM cells in response to TCR stimulation with anti-CD3 + anti-CD28. Floating bars span minimum and maximum, line represents mean. Circles represent individual donors. Wilcoxon matched-pairs signed rank test (a-d), *p < 0.05, ns= not significant.

### TCR-dependent and TCR-independent activation of PLZF^+^ T cells

We next assessed the signals required to trigger cytokine production in mucosal PLZF^+^ T cells and examined their response to stimulation through the T cell receptor (TCR). PLZF^+^ T cells are more responsive to short-term (12-16 hrs) *in vitro* activation by anti-CD3/anti-CD28 monoclonal antibodies (mAb) than PLZF^-^ T cells, as indicated by higher production of both IFNγ and TNFα, which more closely approximated their PMA/Ionomycin induced potential (Fig. 4a, b, Extended Fig. 5a, b). The identification of both IL18R1 and IL18RAP within the core gene signature of PLZF^+^ T cells led us to additionally assess their response to TCR-independent activation. Intestinal PLZF^+^ T cells produced IFNγ in response to the combination of IL-12 and IL-18, but not to either cytokine alone (Fig. 4c). Stimulation with both IL-12 and IL-18 was able to approximate the levels of IFNγ production observed in response to PMA, yet failed to elicit production of TNFα (Extended Fig. 5c). Intestinal PLZF^+^ T cells were more responsive to both TCR- and cytokine-mediated activation than those in the thymus, which failed to produce significant levels of IFNγ and suggested that maturation and acquisition of effector function occurred in the periphery (Extended Fig. 5d). Therefore, we examined whether the memory phenotype of intestinal CD4 T cells was indicative of different functional capacities. We found that the majority of the IFNγ produced by intestinal CD4^+^ T cells in response to either TCR signaling or cytokines alone displayed an effector memory phenotype (PLZF^+^ > PLZF^-^ T cells) (Extended Fig. 5e), and that PLZF^+^ T_EM_ cells produced significantly more IFNγ than PLZF^-^ T_EM_ cells (Fig. 4d). Mucosal PLZF^+^ T cells are therefore poised to generate a pro-inflammatory response to either TCR- and/or cytokine-mediated activation, suggesting that their effector potential may be uniquely adapted for mucosal immune surveillance.

**Figure 5:**
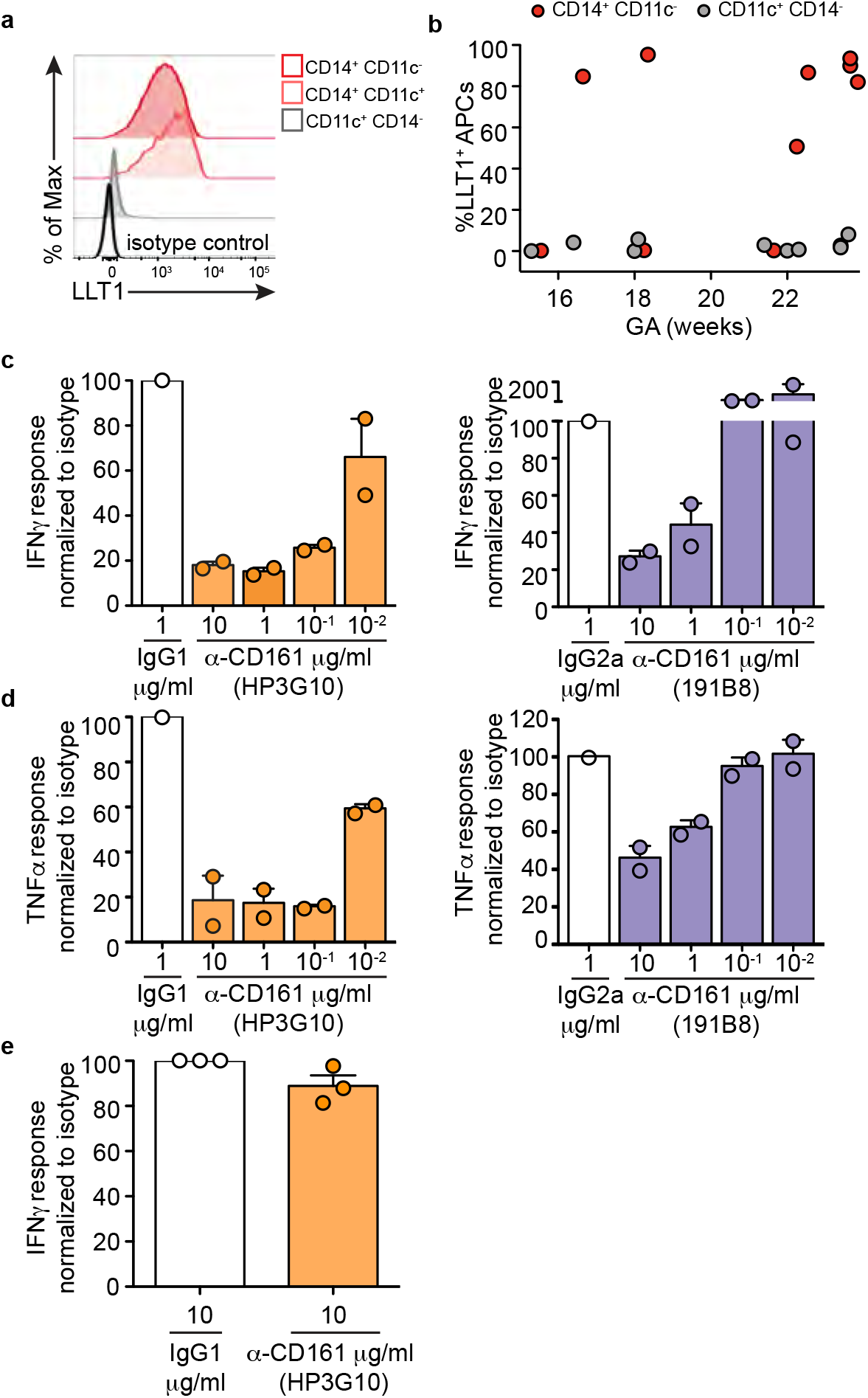
CD161 inhibits production of IFNγ in PLZF+ T cells. a, Histogram of LLT1 expression on indicated intestinal APC subsets. b, Summary of frequencies of LLT1 expression among dendritic cells (CD11c+ CD14-, grey) and macrophages (CD14+ CD11c-, red) of the fetal small intestine relative to gestational age (GA) in weeks. Circles represent individual samples. c, d, Relative proportions of c, IFNγ + and d, TNFα+ PLZF+ Vα7.2-TCRβ+CD4+T cells after 12-14 hrs stimulation with anti-CD3/anti-CD28 in the presence of biotinylated isotypes or anti-CD161 antibodies ligated with anti-biotin beads. One representative experiment of 4 is shown. Bar represents mean with SEM, circles represent technical replicates. e, Relative proportions of IFNγ + PLZF+ Vα7.2-TCRβ+CD4+T cells after 12-14 hrs stimulation with IL-12+IL18 in the presence of biotinylated isotype or anti-CD161 antibodies ligated with anti-biotin beads. Circles represent individual donors. Proportions of IFNγ + and TNFα+ cells are normalized to isotype response. Two different anti-CD161 clones are shown (HP3G10 in orange and 191B8 in purple).

### CD161 inhibits cytokine production in PLZF^+^ T cells

The abundance of polyclonal intestinal (pi) PLZF^+^ T cells capable of responding to two different methods of activation lead us to hypothesize that the behavior of these cells would be tightly regulated. The c-type lectin CD161 is expressed by the majority of intestinal pi-PLZF^+^ T cells and has been ascribed conflicting roles in either the activation or inhibition of different human immune cells ^21,33,34^. LLT1, the natural ligand for CD161, was present on intestinal CD14^+^ antigen presenting cells (APCs), whereas CD11c^+^ CD14^-^ dendritic cells (DCs) lacked expression of LLT1 (Extended Fig. 6a, Fig. 5a). Intestinal CD14^+^ APCs were CD68^+^, CD163^+^, CD209^+^, expressed intermediate levels of HLA-DR, and lacked expression of CD103, consistent with a tissue-resident macrophage phenotype ^35,36^ (Extended Fig. 6b). Macrophage LLT1 expression was not uniform between samples, however LLT1 expression was consistently absent from intestinal DCs (Fig. 5b). The observed variability in LLT1 expression suggested that this ligand was acquired, which prompted us to look for LLT1 expression across tissues with similar populations of APCs. A similar CD14^+^ CD11c^-^ macrophage population in MLN and the appendix mirrored LLT1 expression of intestinal macrophages (Extended Fig. 6c, d), and led us to explore the role of CD161 in the regulation of fetal pi-PLZF^+^ T cells. We found that engagement of CD161 during classical TCR activation resulted in potent inhibition of pi-PLZF^+^ T cells, reducing production of both IFNγ and TNFα (Fig. 5c, d). While two different clones of monoclonal antibodies to CD161 were equally effective at high dose (10μg/mL), clone HP3G10 consistently performed better at lower doses (10^−1^ to 10^−2^μg/mL) (Fig. 5c, d). In contrast, cross-linking of CD161 did not inhibit IFNγ production by PLZF^+^ T cells in response to cytokine stimulation (Fig. 5e).

## DISCUSSION

We have identified a polyclonal effector T cell population with a unique transcriptional profile and Th1 functional properties which is abundant in the human fetal small intestine. The transcriptome of pi-PLZF^+^ T cells displayed an unusual enrichment in myeloid-associated genes compared with conventional CD4 T cells, which predicted the strong effector properties of these cells. In conjunction with the ability to respond to cytokines independently of TCR activation, these gene signatures highlight the innate-like qualities of pi-PLZF^+^ T cells, reminiscent of murine fetal CD5^+^ B cells and TCRγδ T cells ^37–39^, as well as fetal MAIT cells and iNKT cells ^24,27^. Unlike classic innate-like T cells, MsFITs expressed a polyclonal TCR repertoire and accounted for up to half of all CD4 T cells in the intestine. The dominant presence of Th1 pi-PLZF^+^ T cells with a distinct and atypical transcriptome and shared attributes with innate immune cells suggests a role as a first line of defense *in utero* and early life.

*In utero* development presents unique antigenic pressures to the fetus that might drive the development and expansion of pi-PLZF^+^ T cells. Fetal T cells mount an antigen specific response to semi-allogeneic maternal cells present within human fetal tissues, and fetal and neonatal T cells show increased self-reactivity ^3,40–42^. Further, recent reports of bacterial presence in human meconium and amniotic fluid ^43–45^ challenge the paradigm that the fetus develops within a sterile environment *in utero*. Intestinal MsFITs are functionally and anatomically positioned to encounter antigen within swallowed amniotic fluid and could mount an immediate protective response. The development of tissue resident T cell memory is closely coupled to anatomic location ^19^, and is reflected in the increased ability of intestinal pi-PLZF^+^ T cells to produce Th1 cytokines, as well as the preferential expression of the inhibitory receptor CD161. Multiple regulatory transcripts are present in the core gene signature of pi-PLZF^+^ T cells, and we discovered CD161-mediated inhibition pi-PLZF^+^ T cells as a previously unknown mechanism of immune suppression. The presence of LLT1 on intestinal macrophages suggests they likely work in parallel with other regulatory mechanisms to promote tolerance *in utero* ^3,46–48^. We propose that as in the adult, there is regional compartmentalization of the fetal effector response, and that the intestine is a specialized niche which supports the development of protective immunity *in utero*.

## Material and Methods

### Human tissues

All human tissue obtained for analysis in this study was obtained with approval from and under the guidelines of the UCSF Committee on Human Research. Human fetal tissue (small intestine, appendix, mesenteric lymph node (MLN), spleen, liver, and thymus) was obtained under the auspices of UCSF Committee on Human Research (CHR) approved protocols at 15-23 gestational weeks from San Francisco General Hospital after informed consent from elective termination of pregnancy. Samples were excluded in the case of (1) known maternal infection, (2) intrauterine fetal demise, and/or (3) known or suspected chromosomal abnormality. Adult peripheral blood mononuclear cells (PB) derived from TRIMA residues from TRIMA Apheresis collection kits were obtained from healthy donors after informed consent at the Blood Centers of the Pacific. Umbilical cord blood (CB) from term infants was obtained in acid citrate dextrose by sterile cordocentesis from the University of Texas MD Anderson Cancer Center in Houston, Texas.

### Cell Isolation

PBMCs from adult and cord blood were isolated by Ficoll-histopaque (Sigma Aldrich) gradient centrifugation and cryopreserved in freezing medium composed of 90% FBS + 10% DMSO (ATCC). Fetal organs were collected into cold RPMI with 10% FCS, 10 mM HEPES, penicillin, streptomycin, 0.1 mM 2-β-mercaptoethanol, 2 mM L-glutamine, and nonessential amino acids (cgRPMI medium), transported on ice and processed within 2 hours of collection. The small intestine was dissected from the mesentery, cut longitudinally, and meconium was removed with gentle scraping. The intestine was cut into 1cm sections and mucus was removed with three washes in 1mM DTT in PBS for 10 minutes. The epithelial layer was removed with three washes in 1mM EDTA in PBS for 20 minutes. The intestine, MLN, liver, and spleen were digested with freshly prepared 1mg/mL Collagenase IV (Life Technologies) and 10 mg/mL DNAse (Roche) in cgRPMI for 30 minutes, and dissociated cells were filtered through a 70 um strainer. Cells were separated in a 20%-40%-80% Percoll density gradient at 400 × g for 40 minutes. T cells were recovered at the 40-80% interface and APCs were recovered at the 20-40% interface, and all cells washed twice in cgRPMI. All washes and incubations were performed in a shaking (200 rpm) water bath at 37°C. Thymocytes were isolated by gently pressing small pieces of thymus though a 70 um strainer. Viability was measured with Trypan Blue (Sigma Aldrich).

### Antibodies and flow cytometry

Intestinal lamina propria (LP) T cells were isolated by negative selection using the Easy Sep Human T cell isolation kit (STEMCELL Technologies). Isolated cells were incubated in 2% FCS in PBS with 1mM EDTA (staining buffer) with human Fc block (STEMCELL Technologies) and stained with fluorochrome-conjugated antibodies against surface markers. Intracellular protein detection was performed on fixed, permeabilized cells using the Foxp3/Transcription Factor Staining Buffer set (Tonbo Biosciences). Mouse anti-human mAbs used in this study include: TCRβ□DPercp-e710 (Clone IP26, eBioscience Cat. No. 46-9986-42), Vα7.2 Biotin and BV605 (Clone 3C10, BioLegend Cat. No. 351720), CD4 APC H7 (Clone L200, BD Pharmingen Cat. No. 560837), CD8α FITC and PE (Clone B7-1, BD Pharmingen Cat. No. 557226), CD45RA PE (Clone HI100, BD Pharmingen Cat. No. 555489), CCR7 BV421 (Clone G043H7, BioLegend Cat. No. 353208), CD69 PE (Clone FN50, BioLegend Cat. No. 310906), CD103-FITC (Clone Ber-ACT8, BD Pharmingen Cat. No. 550259), PLZF-APC (Clone 6318100, R&D Cat. No. IC2944A), CD161-BV711 (Clone DX12, BD Biosciences Cat. No. 563865), IL18Rα-PE (Clone H44, BioLegend Cat. No. 313808), PD-1 BV605 (Clone EH12.2H7, BioLegend Cat. No. 329924), IFNγ -FITC (Clone 25723.11, BD Biosciences Cat. No. 340449) TNFα-PE Cy7 (Clone MAB11, BD Pharmingen Cat. No. 557647), IL-2 (Clone 5344.11, BD Cat. No. 340448), IL-8 BV421 (Clone G265-8, BD Biosciences Cat. No. 563310), CD45 APC (Clone HI30, Tonbo Cat. No. 20-0459), CD14 BV605 (Clone M5E2, BD Pharmingen Cat. No. 564054), CD11c BB515 (Clone B-ly6, BD Pharmingen Cat. No. 564491), HLA-DR APC-R700 (Clone G46-6, BD Cat. No. 565128), CD3 BV510 (Clone HIT3α, BD Cat. No. 564713), CD19 BV510 (Clone SJ25C1, BD Cat. No. 562947), CD56 BV510 (Clone NCAM16.2, BD Cat. No. 563041), CD68 PE Cy7 (Clone FA-11, BioLegend Cat. No. 137016), CD163 BV711 (Clone GHI/61, BioLegend Cat. No. 333630), CD209 PerCP Cy 5.5 (Clone DCN46, BD Pharmingen Cat. No. 558263), LLT1 PE (Clone 402659 R&D Cat. No. FAB3480P). TCRβ repertoire profiling was performed by staining with the IOTest Beta Mark TCR V β Repertoire Kit (Beckman Coulter). TCRβ□ repertoire profiling was performed by staining with the IOTest Beta Mark TCR V beta Repertoire Kit (Beckman Coulter). All cells were stained with Aqua LIVE/DEAD Fixable Dead Cell Stain Kit (Invitrogen) to exclude dead cells from analysis. All data were acquired with an LSR/Fortessa Dual SORP flow cytometer (BD Biosciences) and analyzed with FlowJo V10.0.8 (TreeStar) software.

### Immunohistochemistry

Intestinal sections were dissected and fixed in 4% paraformaldehyde for 10 minutes and subsequently passed through a sucrose gradient prior to embedding in OCT. Embedded sections were then frozen on dry ice and stored at −80°C. 10 um thin cryosections were obtained using a cryostat and mounted on frosted charged slides. Histological work, imaging, and image processing was performed by the Gladstone Institutes’ Histology & Light Microscopy Core. Slides were fixed in acetone for 5 minutes at −20°C, rehydrated in PBS for 10 minutes, rinsed in 0.05% PBS-Tween, and permeabilized in 0.1% PBS-TritonX-100 for 15 min at room temperature (RT). Slides were then blocked using 1% BSA, 0.1% Fish skin gelatin, 0.5% TritonX-100, 0.05% Na-Azide in PBS for 2 hours and stained with purified anti-CD3 (clone Hit3a; BD Pharmingen, cat. # 555336) and anti-PLZF (clone NBPI-80894, Novus Biologicals, cat. NBPI-80894) overnight at 4°C, followed by anti-CD4-eFluor 570 (N1UG0, Thermo-Fisher, cat. # 41-2444-82) at RT for 3 hours. Slides were then incubated with secondary antibodies goat anti-mouse-DyLight 488 (Thermo Fisher, cat. # 35502) and goat anti-rabbit 633 (Thermo Fisher, cat. # A-21070) for 1 hour at RT, stained with DAPI (Invitrogen) for 10 min at RT, and mounted with Fluormount-G (EMS). Images were captured with a Zeiss Cell Observer Spinning Disk microscope (Carl Zeiss Microscopy, Thornwood, NY) equipped with 405 nm, 488 nm, 561 nm, 633 nm lasers, Prime 95b sCMOS camera (Photometrics, Tucson, AZ) and Zeiss Zen imaging software.

### T cell stimulation and CD161 inhibition

For detection of basal cytokine potential, single cell suspensions from various tissues were cultured directly *ex vivo* in a 96-well U-bottom plate in cgRPMI and stimulated with 50 ng/ml phorbol myristate acetate (PMA) (Santa Cruz Biotechnology) and 5ug/ml ionomycin (Sigma-Aldrich) in the presence of Brefeldin A (Sigma-Aldrich) for 3 hours at 37°C in 4% O_2_ to mimic intra-uterine hypoxia. Alternatively, T cells were cultured at 250 k/well in 96-well plates and stimulated with plate-bound anti-CD3 (clone HIT3a) at 1 ug/ml and soluble anti-CD28 (CD28.2) at 2ug/ml, or cells were activated with IL12 and IL18 (PeproTech) at 50 ng/mL for 12 hours at 37°C in 4% O_2_, and Brefeldin A was added for the last 4 hours. After stimulation, cells were stained for intracellular cytokine production as described above. For the inhibition assays, LP T cells were first incubated with anti-CD161-Biotin mAb (HP3G10, Biolegend or 191B8, Miltenyi Biotec) and isotype controls IgG1 (MPOC-21, BioLegend) and IgG2a (MPOC-173, BioLegend) for 15 minutes, washed, and cross-linked with anti-Biotin Cocktail (STEMCELL Technologies) during stimulation with anti-CD3/anti-CD28 or cytokines as described above.

### T cell isolation and RNA extraction for RNA sequencing

PLZF^+^ and PLZF^-^ CD4^+^ TCRβ^+^ T cells were sorted using a FACS Aria2 SORP (BD Biosciences). T cells were isolated as described above from the intestine of 5 individual samples, and MAIT cells were pooled from the intestine and MLN for RNA sequencing (RNAseq) as outlined in Extended Fig. 2a and 10^4^ cells were collected per subset for each sample. Post-sort purity was determined by flow cytometry following intracellular staining for PLZF as above and was >92% for PLZF^-^ T cells and MAIT cells, and >87% for PLZF^+^ T cells. RNA was extracted and purified with the Dynabeads mRNA DIRECT Purification Kit (Thermo Fisher Scientific). mRNA libraries were constructed using the Nugen/Nextera XT Library Prep Kit (Illumina), and 15 samples (5 donors, 3 T cell subsets) were sequenced on an Illumina HiSeq 4000 by the Functional Genomics Core Facility at the University of California, San Francisco. The reads from the Illumina Hi-seq sequencer in fastq format were verified for quality control using fastqc software package and reads were aligned to the Human genome (hg38) and read counts aggregated by gene using the Ensembl GRCh38.78 annotation using STAR ^49^ Differential gene expression analysis was performed with DESeq2 v1.16.1 package ^50^.

### Statistical analysis

Groups were compared with Prism Version 5 software (GraphPad) using the Wilcoxon matched-pairs signed rank test or the Kruskal Wallis with Dunn’s Multiple comparison test. Box plot rectangles show first to third quartile, the line shows the median and the whiskers represent minimum and maximum values, unless otherwise stated. Correlation analysis was measured by Spearman correlation coefficient. Bar plots represent the mean and the standard error of the mean. p<0.05 was considered significant. Principle component analysis and accompanying confidence intervals were performed using combined functions in the R packages ‘stats’, ‘vegan’, and ‘ggplot2’, and PERMANOVA analysis was used assess the significance of the Euclidean distances between the groups. Heatmaps were generated using the R packages ‘heatmap’ and ‘circlize’. A co-expression analysis was performed for the differentially expressed (DE) genes using the Human Primary Cell Atlas ^29^. This atlas contains 713 microarray samples of a wide range of pure cell types and states. The complexity of the dataset was reduced to the median gene expression per cell state (N=157), and Spearman correlation coefficients were calculated for each pair of DE genes, revealing clusters of genes that are co-expressed in the same set of cell types. Gene clusters were analyzed for overlap with KEGG and GO biological processes gene sets using the Molecular Signatures Database (MSigDB) ^51,52^. No statistical methods were used to predetermine sample size. The experiments were not randomized and the investigators were not blinded to allocation during experiments and outcome assessment.

### Data and software availability

RNA-Seq data that support the findings of this study have been deposited in NCBI BioProject with the primary accession code PRJNA438160. (http://www.ncbi.nlm.nih.gov/bioproject/438160). Further data that support the findings of this study are available from the corresponding authors upon request.

## Acknowledgements

We would like to acknowledge the tissue donors. Thanks to Heather Melichar, Melissa Ng, Adrian Erlebacher, and Tippi Mackenzie for thoughtful critique of this manuscript. UCSF flow core provided instrumentation assistance, which was supported by NIH P30 DK063720 and NIH Shared Instrument Grant 1S10OD021822-01. This study was supported by the UCSF Clinical and Translational Science Institute Pilot Award for Basic and Translational Investigators 2014908. JH was supported by the National Institutes of Health and National Institute of Child Health and Human Development K12 Career Development Award at UCSF K12HD072222 and the National Institutes of Health and National Institute of Allergy and Infectious Diseases K08 Mentored Clinical Scientist Development Award K08 AI128007. ER was supported by National Science Foundation Graduate Research Fellowship and National Institutes of Health and National Institute of Allergy and Infectious Diseases F31 Fellowship 1F31AI136336-01.

## Author contributions

JH conceived of and designed the study, performed research, analyzed the data, and wrote the manuscript; ER contributed to study design, performed research, and manuscript development; DA contributed to bioinformatics analysis and created figures for manuscript; VM performed immune cell isolations; WE contributed to data analysis; TDB contributed to study design. All authors discussed the results and edited the manuscript.

---

**Extended Figure 1:**
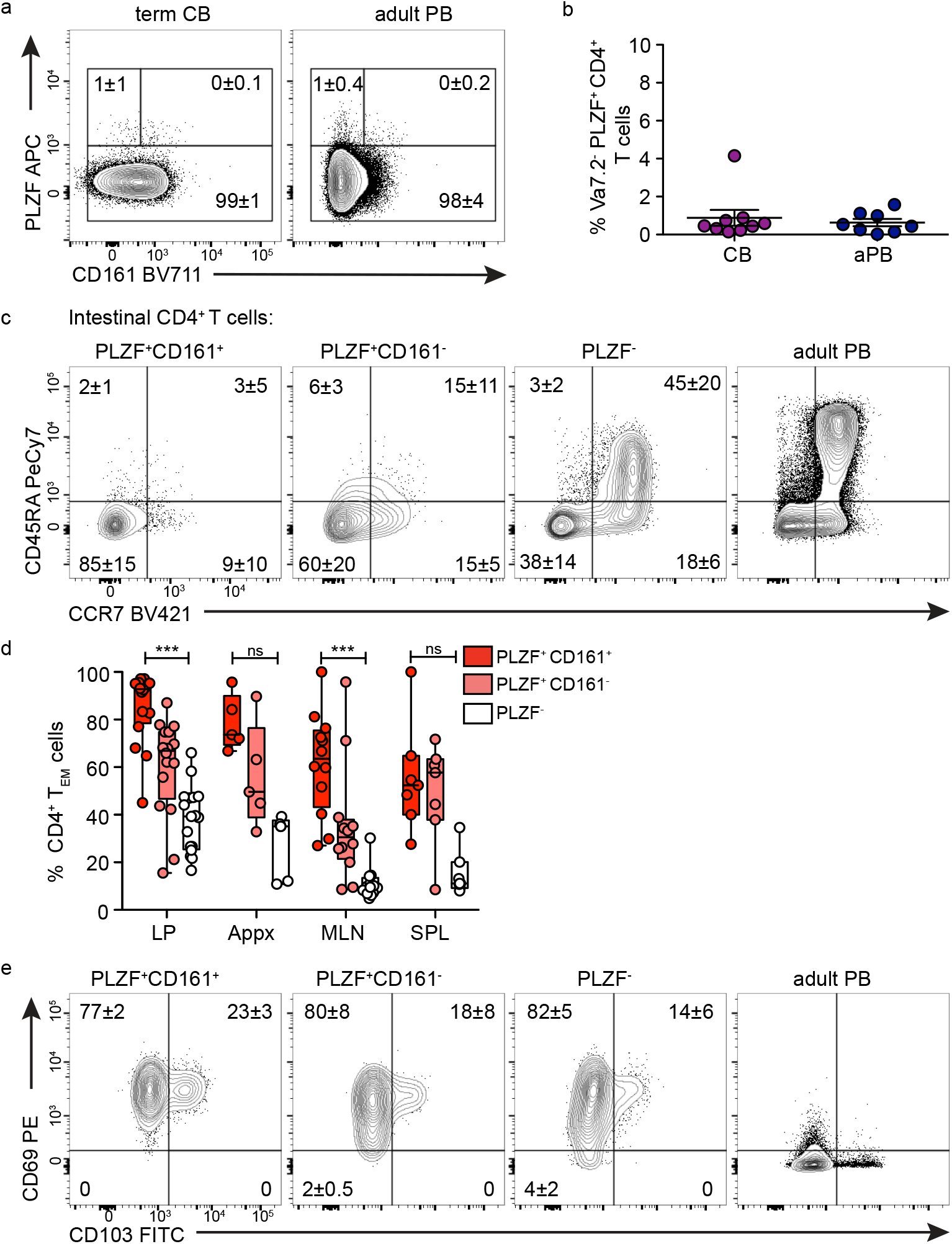
PLZF+ CD4+ T cells are enriched in the fetal small intestine and possess a resident memory phenotype. All samples gated on Vα7.2-TCRβ+CD4+T cells. a, Representative flow plots of PLZF and CD161 expression among CD4+ T cells in term cord blood (CB) and adult peripheral blood (PB). b, Frequencies of PLZF+ CD4+ T cells in term cord blood (CB) and adult peripheral blood (PB). Line represents mean, error bars SEM. c, Representative flow plots of CD45RA and CCR7 expression among indicated subsets of intestinal CD4+ T cells and adult PB gating control. d, Frequencies of CD45RA-CCR7-TEM cells among indicated T cell subsets in fetal tissues. e, Proportions of Naïve, TCM, and TEM within indicated intestinal subsets of CD4+ T cells.e, Representative flow plots of CD69 and CD103 expression among indicated subsets of intestinal CD4+ TEM cells and adult PB gating control. Circles represent individual donors (d). Numbers in the flow cytometry plots correspond to the mean ± SD of gated populations.

**Extended Figure 2:**
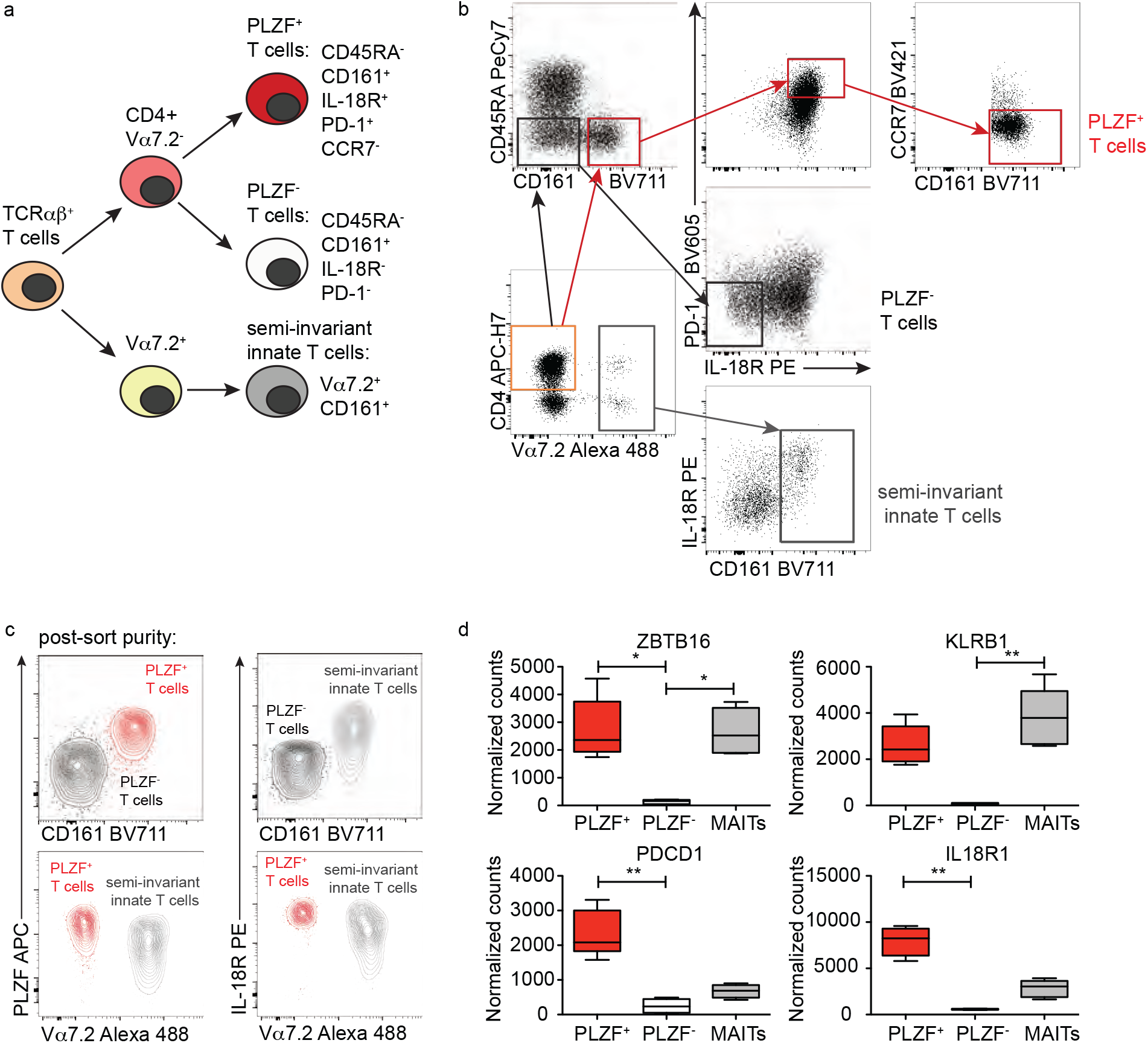
Characteristics of sorted fetal T cell populations. a, Schematic representation of strategy for the identification and isolation of indicated T cell populations. b, Gating strategy for PLZF+ T cells, PLZF- T cells, and MAIT cells after sorting on live, TCRβ+ intestinal T cells. c, Representative flow plots of post-sort purity after intra-nuclear staining for PLZF. d, Boxplots show quantification of RNAseq normalized read counts of indicated genes among sorted populations. *p<0.05, **p<0.01. Kruskal Wallis with Dunn’s Multiple comparison test (d).

**Extended Figure 3:**
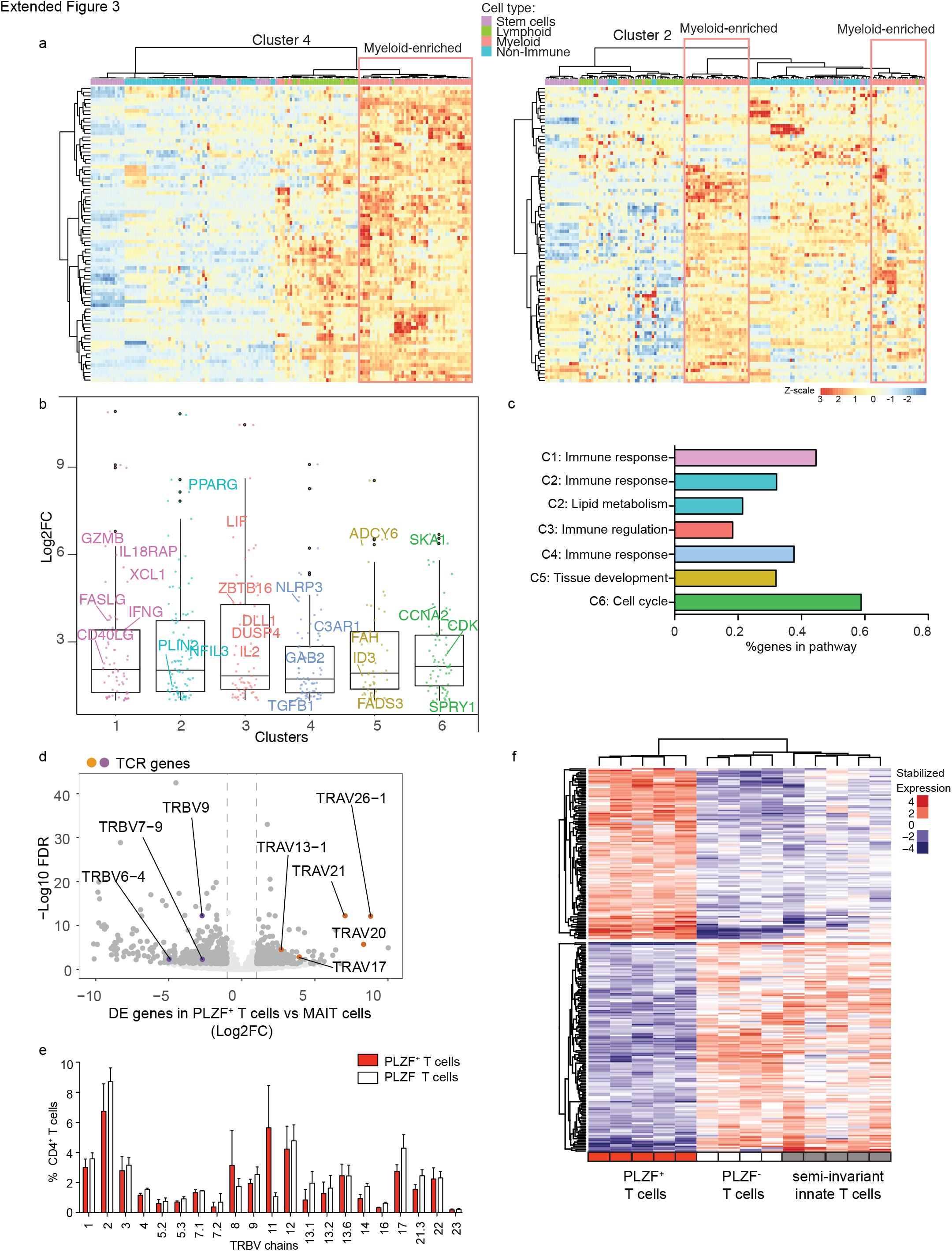
Distinct transcriptional signature of PLZF+ T cells. a, Heatmap shows color-coded relative enrichment of differentially expressed genes in PLZF+ T cells relative to PLZF- T cells among indicated cell types identified from the Human Primary Cell Atlas for Cluster 4 (left) and Cluster 2 (right). b, Comparison of differentially expressed genes in PLZF+ T cells relative to PLZF- T cells identified in Figure 2c. Boxplots show median (centre line), interquartile range (box), 25- and 75 percentiles (whiskers), and outliers (dots above whiskers). c, Pathway analysis of gene clusters identified in Figure 2c was performed and gene pathways were organized into sub-clusters based on shared genes. Bar graphs show the % gene enrichment of each gene pathway within the indicated cluster. d, Volcano plot of differentially expressed genes (>2 fold, FDR<0.05) in PLZF+ T cells compared to MAIT cells. Differentially expressed genes are shown in dark grey, and TCR genes enriched (orange) or depleted (purple) in PLZF+ T cells are labeled. e, Relative levels of expression of 21 TCR Vβ chains on 3 separate intestinal samples. Levels of expression (mean ± SD) in PLZF+ CD4+ T cells (red) are compared to PLZF- CD4+ T cells (white). f, Hierarchical clustering of genes uniquely up- or down-regulated in PLZF+ T cells.

**Extended Figure 4:**
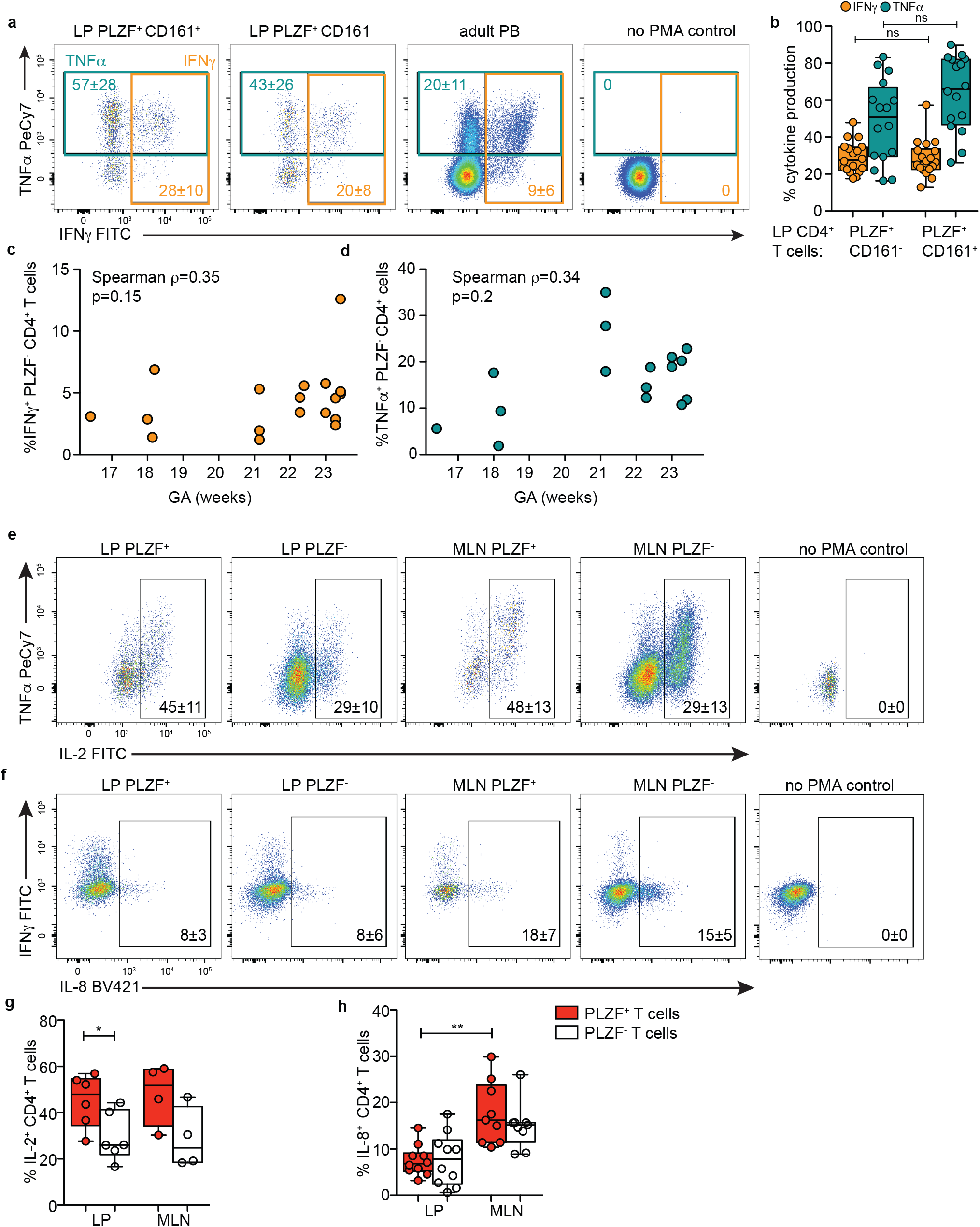
PLZF+ CD4 T cells are poised for rapid production of Th1 cytokines. All samples gated on live Vα7.2-TCRβ+CD4+T cells. a, Representative flow plots of intracellular TNFα and IFNγ staining of indicated populations after PMA/Ionomycin stimulation of intestinal and adult PB CD4 T cells (n=18). b, Frequencies of IFNγ + and TNFα+ cells within indicated populations of CD4 T cells after PMA/Ionomycin stimulation. c, d, Spearman’s rank correlation was used to calculate the association between c, IFNγ and d, TNFα production by indicated subsets of intestinal CD4 T cells in response to PMA/Ionomycin. Circles represent individual donors. e-f, Representative flow plots of intracellular e, IL-2 and f, IL-8 after PMA/Ionomycin stimulation within indicated populations of intestinal and MLN CD4 T cells. g-h, Frequencies of g, IL-2+ and h, IL-8+ cells within indicated populations of CD4 T cells after PMA/Ionomycin stimulation within fetal LP and MLN. Numbers in the flow cytometry plots correspond to the mean ± SD of gated frequencies. Circles represent individual donors. Wilcoxon matched-pairs signed rank test (b, g, h). *p < 0.05, **p < 0.01.

**Extended Figure 5:**
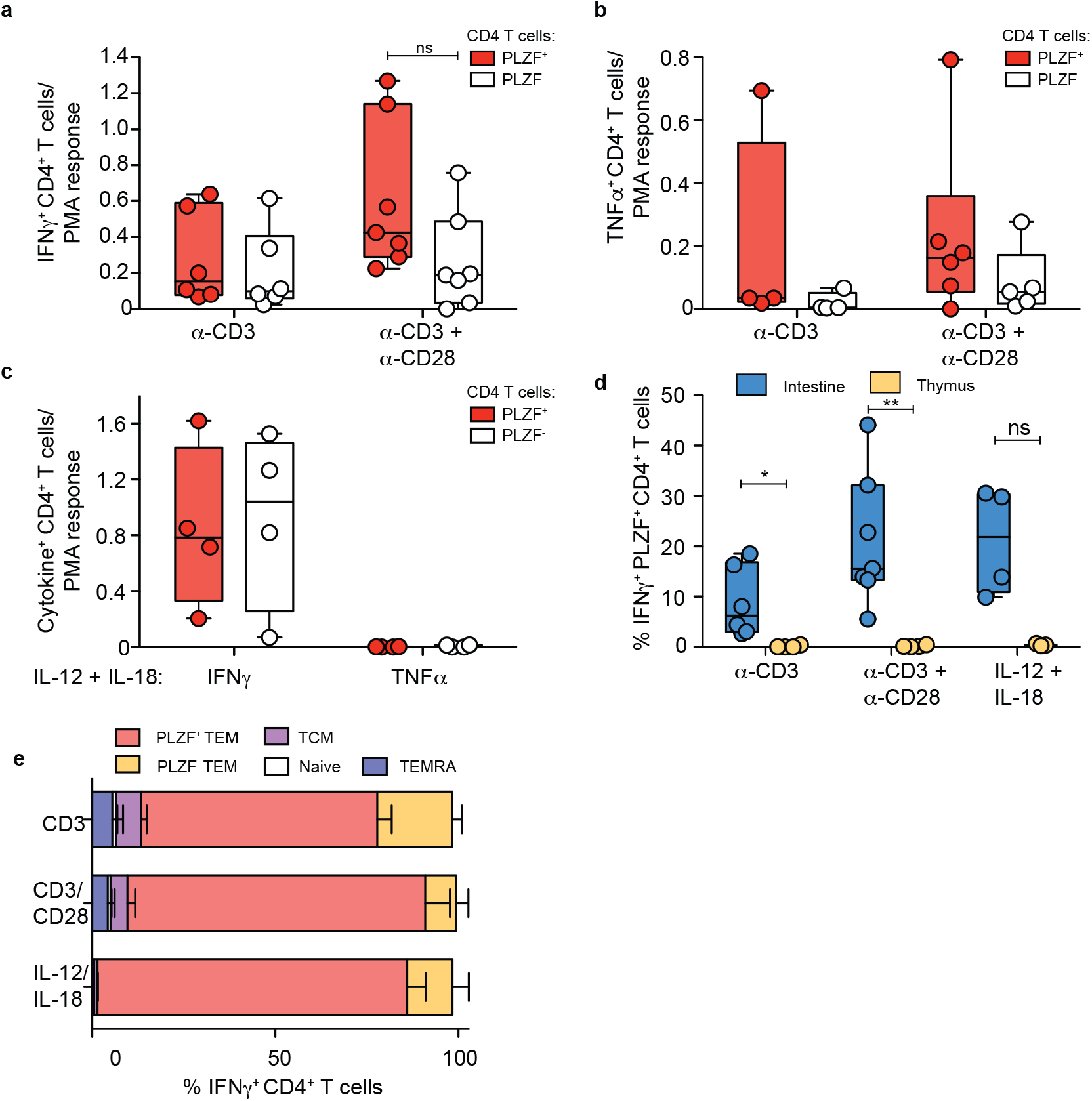
PLZF+ T cells produce Th1 cytokines in response to both TCR- and cytokine-mediated activation. a, b Frequencies of a, IFNγ + and b, TNFα+ cells relative to their PMA/Ionomycin potential within PLZF+ (red) and PLZF-(white) subsets of intestinal Vα7.2-TCRβ+CD4+T cells after 12-14 hrs of TCR stimulation with anti-CD3 +/-CD28. c, Frequencies of IFNγ + cells relative to their PMA/Ionomycin potential within PLZF+ (red) and PLZF-(white) subsets of intestinal Vα7.2-TCRβ+ CD4+T cells after 12-14 hrs of cytokine stimulation with IL-12 and/or IL-18. d, Comparison of intestinal and thymic frequencies of IFNγ + PLZF+ Vα7.2-TCRβ+CD4+T cells after 12-14 hrs of TCR- or cytokine-stimulation as indicated. e, Frequencies of memory and naïve cell subsets among intestinal IFNγ + Vα7.2-TCRβ+CD4+T cells in response to 12-14 hrs of TCR-(n=5) or cytokine-mediated (n=2) stimulation as indicated. Bar graphs denote mean +/-SEM (e). Circles represent individual donors (a-d, f). Wilcoxon matched-pairs signed rank test (a-d, e). *p < 0.05, **p < 0.01.

**Extended Figure 6:**
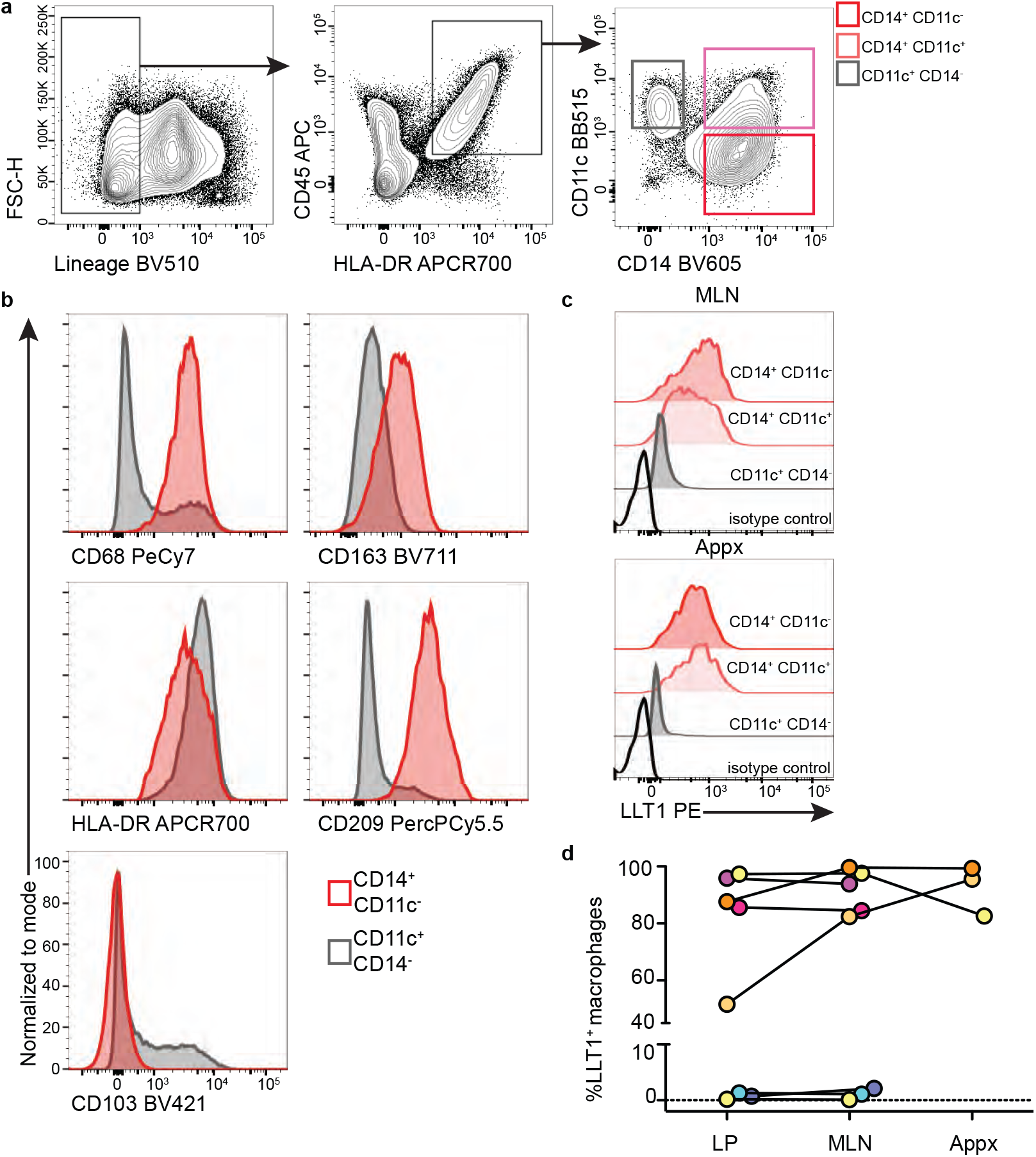
Intestinal macrophages express LLT1, the ligand for CD161. a, Gating strategy for identification of antigen presenting cells (APCs) in fetal tissues. Cells are gated on live, Lineage-(CD3-, CD19-, CD20-, CD56-) intestinal cells and identified as CD45+ HLA-DR+ cells. APCs subsets are defined as CD11c+ CD14-, CD14+ CD11c+, and CD14+ CD11c-. b, Representative histograms of surface markers in CD14+ CD11c-intestinal macrophages as compared to CD11c+ CD14-dendritic cells (n= 12). c, Representative histograms of LLT1 expression on indicated APC subsets in fetal MLN (n=8) and appendix (Appx) (n=3). d, Frequencies of LLT1 expression in tissue macrophages (CD14+ CD11c-) linked by donor. Each color represents an individual donor.

